# Manchester Proteome Profiler: A User-Friendly Platform for Quantitative Proteomic Analysis

**DOI:** 10.64898/2026.05.14.725092

**Authors:** Stuart A. Cain, Mahak Fatima, Martin J. Humphries

## Abstract

Manchester Proteome Profiler (MPP) is an open-source R Shiny application that streamlines downstream analysis of quantitative proteomic data. Compatible with grouped protein intensities tables from MaxQuant, FragPipe, Proteome Discoverer and other custom layouts, MPP provides an integrated platform for filtering, normalisation, imputation, differential expression analysis and cluster analysis across user-chosen experimental conditions. MPP supports both single- and dual-dataset comparisons, incorporates SAINTexpress for affinity purification and proximity labelling experiments, and downstream analysis of the significant protein list clusters to functional enrichment and interaction networks via Gene Ontology, BioGRID and STRING. Benchmarking with a KRAS proximity biotinylation dataset demonstrated the ability of MPP to identify reproducible clusters of differentially expressed proteins and reveal biologically meaningful patterns, including enrichment of solute carrier transporters and adhesion molecules. With interactive visualisations, customisable reports, and support for complex experimental designs, MPP offers a novel, versatile and user-friendly environment for proteomic data exploration and hypothesis generation.

## Introduction

Manchester Proteome Profiler (MPP) (https://mpp.sbs.manchester.ac.uk/) is a user-friendly R shiny application (https://shiny.posit.co/) that has been developed to allow researchers to analyse grouped protein intensity data from MaxQuant ^1^, FragPipe (https://fragpipe.nesvilab.org/), Proteome Discoverer (Thermo Fisher Scientific) or other non-standard protein-intensity tables, that have the same basic format. MPP also has the ability to compare two datasets simultaneously either from different packages or different mass spectrometry techniques (e.g. DDA vs DIA data).

There are several tools available for analysing quantitative proteomic data, which include the R package DEP ^2^ and the R-shiny app LFQ-Analyst ^3^, both of which primarily use MaxQuant data. Perseus, which is companion app to MaxQuant, also offers a comprehensive standalone analysis option, although it relies on manual data handling and is restricted to Windows systems. For analysis of FragPipe protein and peptide intensity data outputs, the R-shiny app FragPipe-Analyst can be used ^4^. MSstats ^5^ can analyse quantitative proteomic data from a number of sources including MaxQuant, FragPipe and Proteome Discoverer, but as an R package requires the user to have some knowledge of R programming. Manchester Proteome Profiler is designed to take proteomic data from several sources, only requires access to a web browser, and allows users to easily query only the condition comparisons they are interested in and to output publication quality figures and processed data files from every step of the data pipeline.

The application has two main sections, the first allows manipulation and quality assessment of the data, and the second performs cluster analysis on the resulting fold-change values generated. Manipulation processes include normalisation and imputation of missing values, while the quality assessments include sample replicate correlation and Principal Component Analysis. The second section of the app allows analysis of the chosen pairwise comparisons, allowing the identification of proteins that have similar log_2_ fold-change profiles across the chosen comparisons, which are then separated into clusters. Clustering information is indicated on volcano plots, and protein intensity information for each cluster can be displayed as grouped box plots. The clusters of significant proteins can be analysed using Gene Ontology to calculate enriched functional profiles using enrichGO, as well as network generation for protein-protein interactions using BioGrid ^6^ or STRING (from known and predicted data) ^7^. All figures generated can be resized and downloaded as PDF files or grouped together as a custom generated report. Manchester Proteome Profiler provides an easy way to identify patterns in the user’s proteomic data to suggest a path for future experimentation.

## Methods

### Application design and data processing pipeline

Manchester Proteome Profiler data analysis is mainly built around three R packages. The first package is the powerful R package, DEP ^2^, which handles filtering, normalisation, imputation, integration of experimental information and quality control. The user inputs their chosen comparisons from their experimental conditions along with their chosen fold-change and adjusted p-value threshold, and the R package ComplexHeatmap ^8, 9^ is used to cluster the resulting significant proteins list. This is then passed onto the third R package, clusterProfiler ^10^, for GO ontology analysis.

The app also allows a significant protein list to be generated using SAINTexpress (Significance Analysis of INTeractome express) ^11^, which can be used for identifying high confidence protein - protein interactions from affinity purification-mass spectrometry (AP-MS) data or proximity biotinylation mass spectrometry. SAINTexpress uses a Bayesian framework to compare protein intensity data from bait samples and negative controls, estimating the probability that a detected prey protein is a true interactor rather than background noise. Samples corresponding to bait samples and prey samples can be identified using the experimental conditions table.

Visualisation of the data is handled by the R packages ggplot2 ^12^, ComplexHeatmap, EnhancedVolcano plots ^13^ and clusterProfiler ^10^, and the reports are built with RMarkdown ^14^ using LaTEX ^15^. The interface is implemented as a Shiny app, using the additional shinywidjets. The full data processing pipeline is summarised in Figure 1.

**Figure 1.**
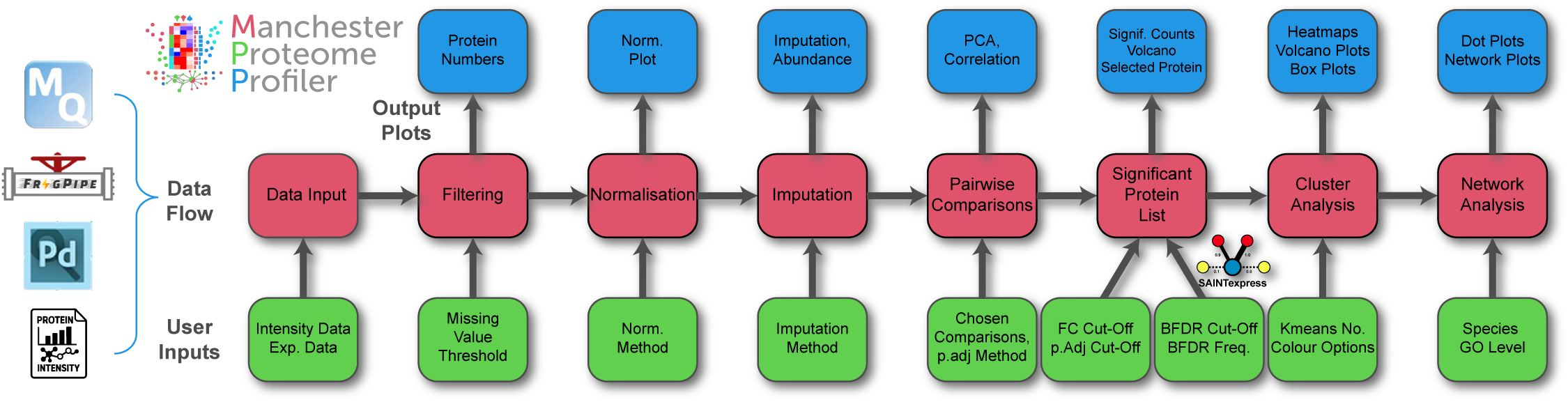
Workflow pipeline for Manchester Proteomic Profile. Users can input protein quantification reports from MaxQuant, FragPipe, Proteome Discoverer or user defined reports from other quantification software such as Spectronaut. This is combined with experimental information (Exp. Data) to initiate the data flow (shown in red). User inputs for each step of the workflow are shown in green and the resultant outputs from each step of the workflow are shown in blu

### Data and condition setup

Manchester Proteome Profiler is designed to process grouped protein intensity files, which can be exported from most quantitative proteomics analysis platforms. These have the same basic layouts, but differ in column annotation, and contain varying amounts of other proteomic information such as peptide counts, etc. There are four standard file input types that can be used with MPP, the *proteinGroups.txt* output from MaxQuant, *combined_proteins.tsv* from the LFQ_MBR workflow of FragPipe, protein group matrix (*report.pg_matrix*) from the DIA_LFQ workflow of FragPipe (or the output from DIA-NN, ^16^) and the “Protein Group Abundance Table” from Proteome Discoverer. The final input option designated “Other” is a flexible non-standard option that will allow any grouped protein intensity file to be imported, if it is in a suitable format. The “Other” data file format allows the user to upload a tab-separated text file, then select the column headings that contain the gene name, accession id, description and amino acid (aa) length. The protein intensity data must be in a continuous block and is identified by selecting the column title of the first and last columns of the block. The accession id, description and aa length columns are optional, as Uniprot data files will attempt to fill in the missing information (human and mouse data). The “Other” option also allows input of Protein Quant Pivot Reports from the DIA proteomics analysis software Spectronaut (Biognosys AG, Switzerland). This output option from Spectronaut has user-selectable additional columns and therefore results in a non-standard format The app also cleans up special characters from the column headers and “NaN” values from the intensity matrix, both of which can cause problems with processing using R.

After the data file has been uploaded, the relevant data columns containing the protein intensity values are extracted depending on the input type: for MaxQuant data, the LFQ_Intensity columns are used and, for FragPipe LFQ_MBR, either the “Intensity” or “MaxLFQ Intensity” columns are used, depending on the radio button selected by the user in the sidebar. FragPipe DIA (or Dia-NN) data input only contains one type of protein intensity data and for Proteome Discoverer the abundance columns of the “Protein Group Abundances” are collected. Columns containing the gene name, protein accession/ID and protein name/description are also extracted. For processing with SAINTexpress, columns containing the protein length are also extracted; however, for DIA data, this data has to be extracted from UniProt data files, which must be either Human or Mouse.

To allow differential expression analysis to be performed, basic experimental metadata is needed to match the samples/columns from the data files with their corresponding experimental conditions. This is achieved by creating a SummarizedExperiment object, a S4 class container ^17^ that merges the protein intensity data with an experimental template file. To assist the user, and ensure the correct intensity columns are matched, a customised experimental template can be downloaded, which contains five columns (label, condition, replicate, condition_type and bait_id). The label column is automatically filled in from the column titles from the intensity data columns, the user then fills in the condition column to group the samples to their chosen experimental name or variable, the replicate number is generated from the last character of the column titles but should be altered if not correct. The final two columns are required for SAINTexpress analysis. SAINTexpress identifies proteins that are significantly enriched relative to their corresponding intensities in control samples. The bait_id column must be annotated with “T” for test samples and “C” for control samples. When multiple sets of controls correspond to specific experimental groups, these can be assigned unique identifiers in the condition_type column, allowing separate SAINTexpress analyses. This column accepts any alphanumeric value and is also used to segment samples in the PCA plot. If SAINTexpress analysis is not performed, these columns can remain unchanged.

MPP uses a pairwise comparison list generated by the user. This allows a significant protein list to be generated that is more specific to the user’s experimental reasoning, rather than comparing conditions that have no experimental relevance. The pairwise comparisons can either be generated from the drop-down list on the sidebar, which lists all possible combination pairs from the experimental template table. However, if the experiment consists of more than a few conditions, the pairwise comparisons can be specified by uploading a comparison table. To facilitate this, a custom comparison template can be downloaded, which already lists the conditions from the experimental template file. This allows the user to quickly copy and paste the comparisons into the relevant conditions into “Comparison A” and “Comparisons B” columns. Once uploaded, the columns are then converted into pairwise queries “Comparison A” vs “Comparisons B”.

The app includes an option to perform two sets of comparisons using distinct pairwise conditions and separate cut-off values. This generates two independent lists of significant proteins, which are then intersected to identify proteins common to both sets—forming the final significant protein list. This approach enables more complex queries, such as identifying proteins that are significantly different between drug-treated and control conditions, and the ability to show significant differences across multiple time points.

### Two dataset comparison mode

If the Two-Dataset comparison mode toggle is activated, this now allows two separate data files to be loaded. These can be from the same datatype or different (e.g. MaxQuant and FragPipe analysis, or FragPipe DDA and DIA data). Each must have is own experimental template file, which can be generated from its dataset file. The comparison template file will now have a sixth column containing the list of the conditions from the second experimental template, which must be used to generate the conditions for the second round of comparisons (4th and 5th columns).

### Filtering, normalisation and imputation

Proteins with excessive missing values are removed prior to analysis. The default filter threshold of 1 excludes proteins lacking intensity values in all but one replicate within any condition, whereas a threshold of 0 (most stringent) retains only proteins quantified in all replicates of at least one condition. The data can then normalised, the default method being variance stabilising transformation (VSN), which minimises the dependence of variance on mean intensity and improves comparability across samples. The normalisation method can be adjusted via the “Normalisation options” dropdown menu in the sidebar, allowing users to choose between median-centred, quantile, or no normalisation. When the two-dataset mode is enabled, different normalisation methods can be applied to each dataset independently, facilitating direct comparison of their outputs.

Imputation of missing values has a default method (“man”) where missing values are generated using random draws from a manually defined left-shifted Gaussian distribution centred around a minimal value (used for missing not at random data (MNAR)). Other imputation options can also be selected using the radio buttons. These include methods that use imputation methods employing stochastic elements, so can differ slightly with different random seeds. These methods include the default method “man”, QRILC (Quantile Regression Imputation of Left-Censored data) and “MinProb” (Minimal value with added noise). To overcome this problem, it is possible to impute the missing values with three different random seeds and then use the average imputed value. The original non-missing values remained unchanged. Users can select this option from the “Imputation options” menu. Other methods are available that are not affected by random seed values, such as Nearest Neighbour Averaging (knn), Bayesian Missing Value Imputation (bpca) or Nonlinear Iterative Partial Least Squares (NIPALS). The application also implements a custom hybrid imputation method, “knn + man”, which combines k-nearest neighbour and manual (minimum-based) approaches. In this method, the normalised data is first divided by experimental conditions. For each condition, missing values are imputed using k-nearest neighbour estimation when at least two replicates contain valid intensity measurements. Remaining missing values, those occurring in conditions with one or no valid replicates, are subsequently imputed using the default “man” method. As with normalisation, different imputation strategies can be applied in the two-dataset mode, enabling users to compare the impact of different imputation settings between datasets.

### Generating significant protein list

Once missing values have been imputed, differential expression analysis can be performed across the entire dataset using linear models implemented via the *limma* R package ^18^. Users provide a contrast matrix, enabling the definition of complex comparisons beyond simple control-versus-treatment setups or exhaustive pairwise combinations. Fold-change and adjusted p-value (adj .p) threshold values can be set using input boxes at the top of the app and can be set to two independent values if two comparison or two-dataset methods are used. The default method for generating adjusted p-values uses the *fdrtool* ^19^ for estimating both tail area-based false discovery rates (Fdr). This can be changed to other methods such as “BH” (Benjamini-Hochberg) or “Bonferroni”, using the side bar menu. Like the normalisation and imputation methods, different options can be selected in the two-dataset mode to allow the user the compare the outputs.

### Cluster analysis

After a significant protein list is generated, a heatmap is generated using the list displaying log_2_ fold-change values from the chosen comparisons. The heatmap is generated using the ComplexHeatmap package ^9^ and includes violin plots to show the distribution of the log_2_FC values for each comparison. A single row heatmap shows the mean log_2_Fc for each comparison and a horizontal histogram shows the number of times each protein has passed the significant threshold (which will range from one to the total number of comparisons). K-means clustering is used to divide the significant protein list into different cluster groups. This helps the user to identify patterns and structures within the data and is also used to set the colour of the points in the EnhancedVolcano and network plots, and to help the user identify how the clusters relate to each comparison. Box plots showing the log_2_ intensity values of both the imputed and non-imputed (but normalised) data and can also be seen for each of the cluster groups. The box plots also show each individual replicate points. A separate complex heatmap is also generated for each cluster, which includes violin plots and mean log_2_FC specific for that cluster group. This allows easier representation of the data in a publication if the user is describing the data for a specific cluster.

### Gene Ontology and protein-protein network analysis

The cluster groups generated from the heatmap are subsequently used for gene cluster enrichment analysis with the clusterProfiler package ^10^, enabling the identification of overrepresented biological processes. Both dot plots and Concept Network plots (cnetplot) are produced for the three primary Gene Ontology categories: Molecular Function, Cellular Component and Biological Process. The hierarchical level of each plot can be adjusted to display either broader, more general terms or more specific, detailed categories.

BioGrid is a biomedical interaction repository (https://thebiogrid.org/) ^6^ from which a network plot can be generated by performing a search using the significant protein list in the biogridr package ^20^. The search output is then filtered so it solely contains interactions between proteins on the list. Network complexity can be controlled by setting the minimum times an interaction entry between a pair of proteins is listed. This is controlled by the “Interaction Count” input box and is also indicated by the thickness of the line between the two proteins. The BioGrid database has 10-fold more interactions for human proteins than mouse, so if a more detailed network is needed with mouse protein data, the names can be converted to uppercase and searched against the human interaction network using the “Use Human Interaction Network Data” checkbox. This option is not suitable for Arabidopsis or Xenopus data.

STRING (Search Tool for the Retrieval of Interacting Genes/Proteins) provides known and predicted protein-protein interactions (https://string-db.org) and is queried using the STRINGdb package ^7^. Interactions between proteins are given a confidence score ranging from 0 to 1, with 1 representing the highest level of confidence. The level of confidence can be increased from the default value of 0.4 to filter out interactions of lower confidence, and the confidence score is represented by the intensity of the line between the two proteins. Node sizes of both plots can be controlled by the “Node Size” input box.

### Report generation and saving results

Every plot generated by the app can be downloaded as a PDF file, with inputs provided to alter the plot size, so an optimum scaled plot can be generated. Tab-delimited text files can also be downloaded containing the processed data from each step of the data pipeline, starting from the normalised filtered data and ending with data tables used to generate the Gene Ontology plots.

Extensive reports can also be generated using Rmarkdown scripts, which include a context-driven method description, a summary of the input variables chosen, including comparisons, and a summary of the number of significant proteins. This is followed by all the plots generated by the app, including tables listing the proteins, their description and accession id for each cluster. As generating the Gene Ontology plots can take several minutes, a “short” report (which can typically be around 35 pages), containing all the plots apart from the GO plots and box plots, or a short report with box plots can also be generated.

### Dataset and processing

The datasets used in this manuscript were generated from the .Raw files available from the PRIDE database (https://www.ebi.ac.uk/pride/) under the PXD069779 identifier. For this study, peptide samples were subjected to liquid chromatography-MS (LC-MS) using a Thermo Q-Exactive HF mass spectrometer (ThermoFisher). The resulting 18 raw files were processed with both MaxQuant and FragPipe. The raw mass spectrometry data were processed using MaxQuant (version 2.1.4.0) with a 1% false discovery rate (FDR) applied at the PSM, protein, and site levels. Label-free quantification was performed with ‘match between runs’ enabled, and protein quantification was based on razor peptides with modifications including Oxidation (M), Acetyl (Protein N-term), and Carbamidomethyl (C). The subsequent grouped protein file (*proteinGroups.txt*), which summarises identified protein groups, their quantitative values, and associated metadata, including accession numbers, peptide counts, modification information, and LFQ intensities and was then used as the data input for the MPP app. Raw MS data was also analysed with FragPipe (version 23.1) using the default ‘LFQ-MBR’ workflow, which integrates MSFragger for peptide-spectrum matching, Philosopher for validation and protein inference with a 1% FDR at PSM, peptide, and protein levels, and IonQuant for label-free quantification with match-between-runs alignment. The resultant *combined_protein.tsv* file which contains the final list of identified protein groups, along with associated information such as protein accessions, gene names, sequence coverage, peptide counts (unique and total) and quantitative values including label-free quantification (LFQ) intensities which was used as the input data for the app.

## Results and discussion

The exemplar dataset used was from a proximity biotinylation (BioID) experiment using a single bait protein (BirA-KRas) and control bait (BirA), which was carried out in 3 separate cell lines. The cell lines were human pancreatic cancer cell lines KP4, which expresses KRAS^G12D^ ^21^ and Suit-2, which expresses KRAS^G12D^, CDKN2A^H83Y^ and TP53^R273H 22^. The third cell line was H6c7, which is derived from normal human pancreatic ductal epithelium and expresses wild-type KRAS ^23^. The BioID experiment was repeated 3 times, which therefore resulted in a total of 18 samples. To demonstrate the capabilities of the Manchester Proteome Profiler app, two analyses were performed. The first shows the results of two comparison rounds: the first round compares the BirA-KRAS against the corresponding BirA only control for each cell line. The second comparison compares the BirA-KRAS protein intensities between the three cell lines. The second analysis uses the two-dataset method and compares the BirA-KRAS against the corresponding BirA control for each cell line using the output from MaxQuant and FragPipe.

### Two-comparison analysis

The *proteinGroups.txt* file from MaxQuant was uploaded into the app and a data-specific experimental template was generated using the columns containing the LFQ intensities. The condition and replicates columns were filled in with the six conditions, and there were three replicates for each. The condition_type column was used to indicate the cell line, and the bait_id was filled in with “C” for the control samples and “T” for the KRAS samples. The last two columns are only needed if SAINTexpress analysis is needed, although the PCA plot can be split into groups using the condition_type information. After the edited experimental template was uploaded, a comparison template was generated, and two sets of comparisons were added. The first round of pairwise comparisons, which compared bait vs control, was KRas_KP4 vs Con_KP4, KRas_Suit2 vs Con_Suit2 and KRas_h6c7 vs Con_h6c7 and the second round of pairwise comparisons, which compared KRAS samples between cell lines was KRas_KP4 vs KRas_Suit2, KRas_KP4 vs KRas_h6c7 and KRas_h6c7 vs KRas_Suit2. The edited comparison file was uploaded, and the analysis was run. The output from MaxQuant contained 2795 proteins groups, of which 1939 proteins were reproducibly quantified after filtering with a missing value set to 1 (for a protein to be included, it had to be present in two or more replicates for at least one of the conditions). The default normalisation method used was variance stabilisation normalisation (VSN), combined with the default “man” imputation approach and the tail area-based false discovery rate (FDR) method for adjusted p-value calculation. Principal Component Analysis (PCA) of the top 500 most variable proteins across samples showed tight clustering of replicates within the conditions, indicating good reproducibility (Supporting Figure S1A), this was also confirmed by the Pearson correlation plot (Supporting Figure S1C). The PCA Scree Plot indicated that the first two principal components accounted for over 50% of the total variance (Supporting Figure S1B). Other Principal Components can be plotted using the app. The Pearson correlation plot also showed clear separation between the KRAS conditions and the control groups.

Based on previous analysis ^24^ and after inspection of initial volcano plots from both sets of comparisons, displayed in the “Result Plots” section of the app (Supporting Figure S1D and S1E), both fold-change cut-off values were set to >= 10 (Log_2_ value 3.32) for each comparison, with identical adjusted p-value cut-off values of <= 0.05. After two sets of pairwise comparisons were performed, 192 proteins differed significantly for the first set of comparisons (bait vs control) (Supporting Figure S2A), and 146 proteins differed in the second set of comparisons (KRAS vs KRAS from each cell line) (Supporting Figure S2B). There were 52 proteins that overlapped the two sets of comparisons, which is indicated by the Venn diagram (Supporting Figure S2C). These proteins form the significant protein list which is used for visualisation and analysis.

### Cluster Analysis

The second part of the Manchester Proteomic Profiler focusses on clustering of the significant protein list. This is achieved using K-means clustering of the log₂ FC values of the chosen comparisons and is displayed in the form of a heatmap (Figure 2A). By using the heatmap visualisation, the appropriate cluster number could be chosen. The 52 proteins that overlapped the two sets of comparisons were separated into four distinct clusters. Using the heatmap and the corresponding volcano plots (Figure 2B–D), each cluster could be broadly classified. Cluster 1 contained proteins with higher detected intensities in the Suit-2 cell line. Cluster 2 included proteins more abundant in the H6c7 cell line. Cluster 3 featured proteins with elevated intensities in KP4 cells, particularly when compared to H6c7. Finally, Cluster 4 comprised proteins that were generally more abundant in KP4 cells, especially relative to Suit-2.

**Figure 2.**
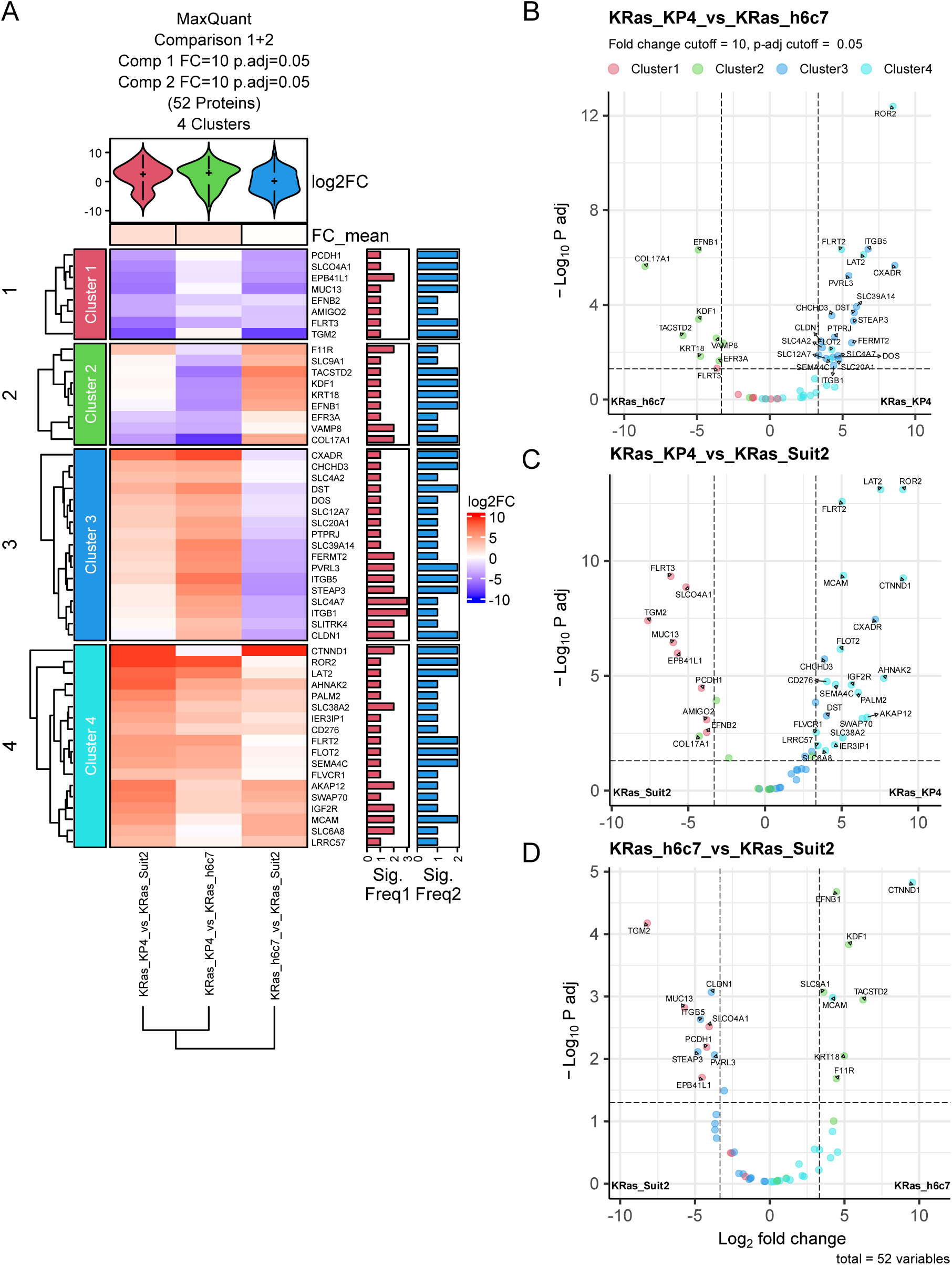
Heatmap and volcano plots generated by Manchester Proteome Profile. **(A)** Heatmap displays log₂ fold-changes of 52 differentially expressed proteins (FC ≥ 10, adj. p ≤ 0.05) that were significant for both sets of comparisons KRas_KP4 vs Con_KP4, KRas_Suit2 vs Con_Suit2, KRas_h6c7 vs Con_h6c7 (Supplementary Figure S2A). and the second round of pairwise comparisons, KRas_KP4 vs KRas_Suit2, KRas_KP4 vs KRas_h6c7, KRas_h6c7 vs KRas_Suit2 (Supplementary Figure S2B).K-means clustering grouped proteins into four clusters, shown left. The violin plot summarises the distribution of fold-changes for each comparison and the bar plots to the right indicate the number of times the corresponding protein was significant for the first round of comparisons (Sig. Freq 1) and the second round of the comparisons (Sig. Freq 2). Volcano plots of comparisons **(B)** KRas_KP4 vs KRas_Suit2, **(C)** KRas_KP4 vs KRas_h6c7, **(D)** KRas_h6c7 vs KRas_ Suit2 indicating the 52 significant proteins for each of the 2 cutoff values (indicated by dotted line). Significant proteins are labelled and points are coloured by the cluster group indicated in the heatmap.

The significant protein list and cluster information was then used for Gene Ontology, BioGRID and STRING analysis. GO Cellular Component enrichment analysis showed enrichment for focal adhesion and cell-substrate junctions in Clusters 1, 3 and 4 (Cluster 2 had no Cellular Component enrichment.) with Cluster 3 enriched for plasma membrane protein and Cluster 4 coated vesicles (Figure 3A). Cluster 3 and 4 contained proteins that were of higher intensity in KP4 cells, and the specific proteins included focal adhesion/lamellipodium components integrin β1 (ITGB1) and kindlin-2 (FERMT2), as well as various transporter proteins (SLC4A7, SLC39A14, SLC4A2). The Category-gene network plot (cnetplot) showing the individual proteins for each enriched term is shown in Figure 3B. Cluster 1, 3 and 4 showed enrichment for Molecular Function terms (Supporting Figure S3A), with most of the proteins involved in the enrichment terms being the ones with higher intensities in KP4 cells. As well as integrin-related proteins (ITGB1, ITGB5 and FERMT2), there are seven symporter proteins, three of which were the same as identified Cellular Component proteins (SLC4A7, SLC39A14, SLC4A2), but also additionally SLC20A1, SLC12A7, SLC38A2, SLC6A8 in Cluster 3 and 4, with SLC04A1 enriched in Cluster 1 (Supporting Figure S3B). Biological Process GO term analysis also indicated enrichment of proteins in Cluster 1 and 2, which were shown to be of a higher intensity in the Suit-2 and H6c7 cell lines, respectively. Cluster 1 proteins AMIGO2, FLRT2, TGM2 and PCDH1, mapped to GO terms cell-cell adhesion, while Cluster 2 proteins mapped to protein localisation and stress fibre assembly. (Figures 3C and 3D). The BioGRID analysis shows proteins mainly from cluster 3 and 4 are centred around two proteins MCAM (Melanoma Cell Adhesion Molecule) and CXADR (Coxsackievirus and Adenovirus Receptor). Both proteins are involved in cell adhesion, and have interactions listed in the BioGrid with the symporters SLC38A2, SLC6A8, SLC9A1, SLC20A1, SLC4A7, FLVCR1, STEAP3. (Supporting Figure S3C). The STRING network was simpler, showing interactions mainly around ITGB1, although the symporters were clustered in two groups (Supporting Figure S3D).

**Figure 3.**
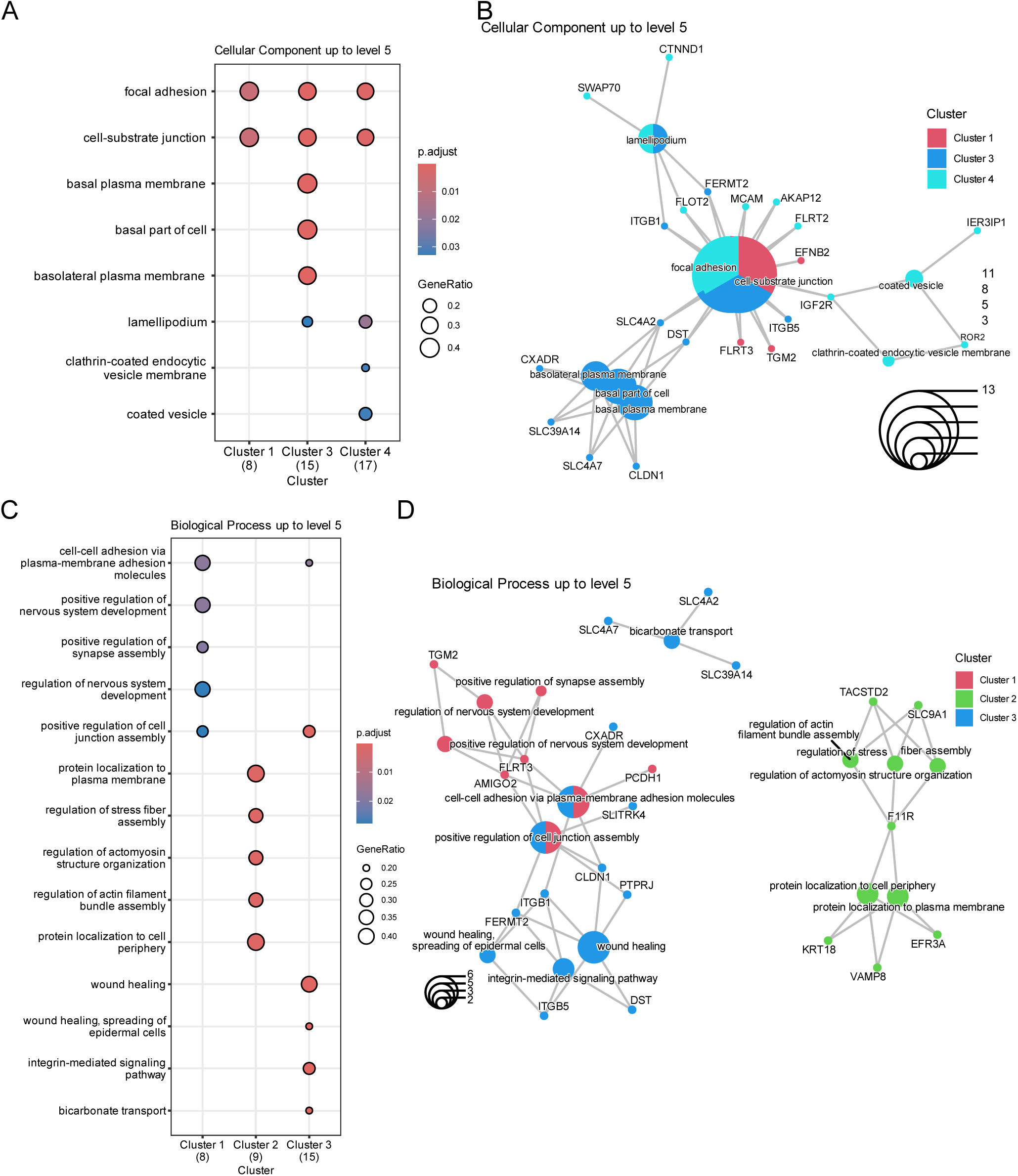
Gene Ontology plots of significant proteins generated by Manchester Proteome Profile. **(A)** Dot plot shows enriched Cellular Component GO terms in Clusters 1, 3 and 4. Cluster 2 showed no enriched GO terms. **(B)** Corresponding proteins from each Cellular Component GO term shown as a category-gene network plot (cnetplot). Node colours correspond to the same cluster colours as the heatmap, and will change if a new palette is chosen for the original heatmap. **(C)** Dot plot shows enriched Biological Process GO terms in Clusters 1, 2 and 3. Cluster 4 showed no enriched GO terms. **(D)** Corresponding proteins from each Biological Process GO term shown as a category-gene network plot.

### Result validation

It is important once cluster analysis has been undertaken that the fold-change differences are not due to artefacts caused by incorrectly assigned imputation values. The best way to check this is to go back to the original normalised intensity values, which are shown in the box plots section. The box plots for the significant protein list are separated into their respective cluster numbers, and box plots can either show the normalised values, with gaps where no protein intensities were available or the imputed values which has the missing replicate protein intensities filled in.

Cluster 3 showed proteins with higher detected intensities in the KP4 cell line, especially compared with H6c7. This can be seen in the individual cluster heatmap for Cluster 3 (Figure 4A), where the violin plot shows a higher log₂ FC values for KRas_KP4 vs KRas_h6c7 and for KRas_KP4 vs KRas_Suit2 (Figures 2B-C). There is also an indication that proteins in this cluster have a lower intensity in H6c7 calls than Suit2 cells. When the Cluster 3 box plots are inspected, it can indeed be seen that all the proteins have a higher intensity in the KRas_KP4 samples. (Figure 4B). There are also high intensity values seen in the KRas_Suit2 samples, although usually lower than the KRas_KP4 samples, apart from CHCHD3 and CXADR. It can also be seen that the protein intensities for the KRas_h6c7 samples are lower or missing completely, indicating these proteins are less abundant in this cell line. When the imputed data for Cluster 3 is inspected (Supporting Figure S4A), it can be seen the imputed values do not cause artefacts giving rise to anomalous log₂ FC values.

**Figure 4.**
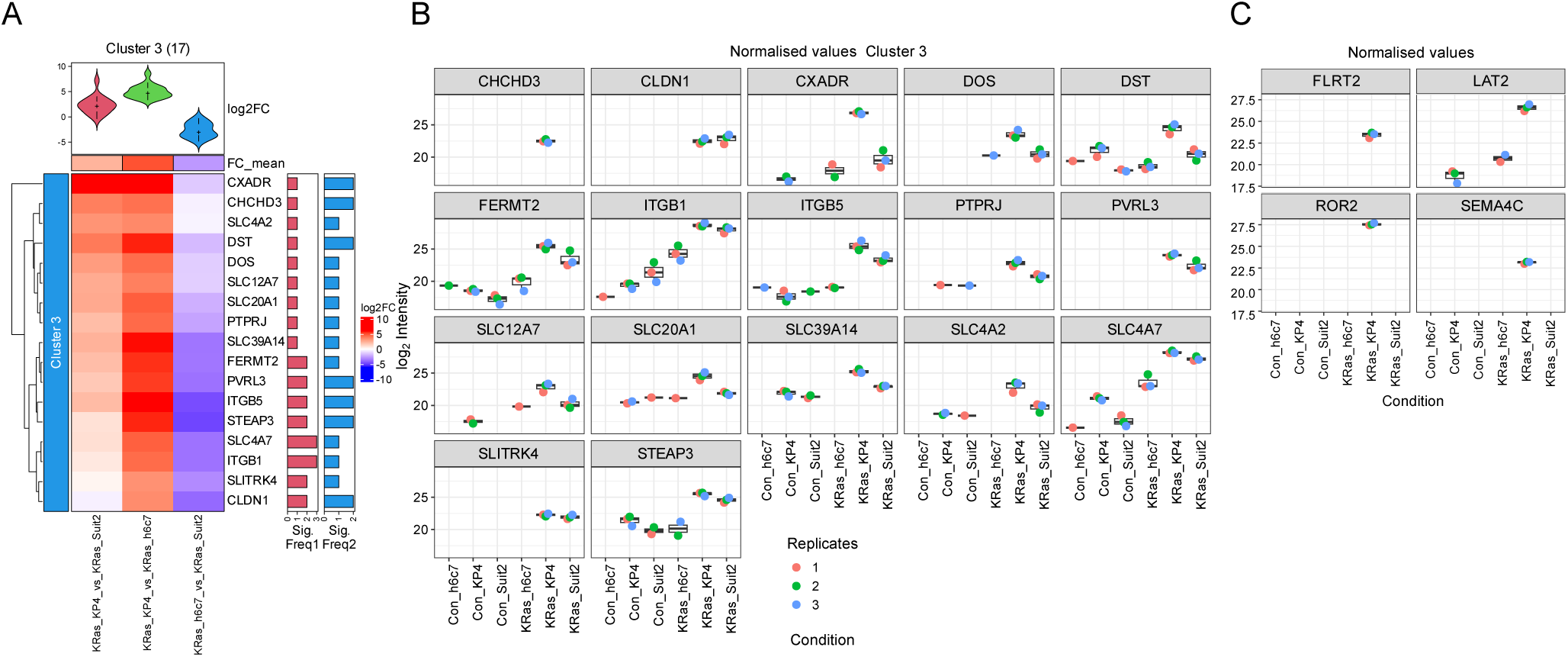
Single cluster heatmap and box plots. **(A)** Heatmap of significant proteins in Cluster 3. The violin plots now summarise the distribution of fold changes for each comparison for that particular cluster. **(B)** Box plots of the normalised protein intensities before imputation of missing values. The box plots provide an easy way to validate the significant protein clusters original data to allow potential target validation by other experimental means. **(C)** Box plots showing normalised values of 4 key proteins that are uniquely or highly enriched in KRas_KP4 samples from Cluster 4. Custom box plot panels can be created in the app on the initial “DEP”tab.

Cluster 2 showed proteins with greater intensities in hc67 cell line, which is indicated with the corresponding violin plots shown in the Cluster 2 heatmap (Supporting Figure S4B). Inspection of the normalised data shows higher intensities or intensities that are on seen in KRas_hc67 samples (Supporting Figure S4C). The protein KRT18, however, can be discounted as it has a higher intensity in the control hc67 sample (Con_h6c7). Again, no anomalous imputed values are seen.

Cluster 4 also contains proteins that are significantly higher in KRas_KP4 samples, especially compared to KRas_Suit2 samples (Supporting Figure S4D-E), some of which such as FLRT2, ROR2 and SEMA4C are only exclusively found in KRAS_KP4 samples. These proteins are not seen in the control sample for KP4, suggesting that these proteins are in proximity to KRAS in KP4 cells (Figure 4C).

### Conclusion for cell comparison analysis

The 52 proteins in the protein list were the overlap between proteins that were of higher intensity than their respective BirA only controls, and protein intensities that were different between the KRAS samples from the different cell lines. The proteins clustered into 4 distinct clusters, of which the bulk of the proteins were in clusters 3 and 4, which correspond to proteins with higher intensities in KP4 cells. These proteins included adhesion molecules such as integrins (ITGB1 and ITGB5), the adhesion adaptor kindlin-2 (FERMT2), and symporters SLC38A2 (SNAT2) and SLC6A8. It has been found that oncogenic KRAS mutations such as G12D upregulate the amino acid transporter SLC38A2 (SNAT2) via YAP1 activation, enhancing glutamine uptake and mTOR signalling to support tumour proliferation ^25^. The creatine transporter SLC6A8 is also reported upregulated in KRAS mutant colorectal cancer cells and to support hypoxic survival ^26^. Cluster 4 includes Melanoma Cell Adhesion Molecule (MCAM), which in BioGRID is a key hub that connects adhesion proteins (CTNND1, FLRT2), signalling molecules (ROR2, SEMA4C, IGF2R, CD276), and transporters (SLC38A2, SLC6A8, FLVCR1 and LAT2 (SLC7A8)). Of the 4 proteins that are either only seen in KRas_KP4 samples (FLRT2, ROR2, SEMA4C), or highly enriched in KP4 cells (LAT2), all four have either established links to oncogenic KRAS biology or tumour aggressiveness. ROR2 has been identified as a regulator of cellular plasticity in pancreatic neoplasia, where it shown to promote epithelial-mesenchymal transition, which can also be hijacked by cancer cells to become invasive and metastatic. ROR2 drives an aggressive PDAC phenotype and confers resistance to KRAS inhibitors ^27^. LAT2 (SLC7A8) promotes glutamine-dependent mTOR activation and glycolysis in pancreatic cancer, contributing to gemcitabine resistance and poor prognosis ^28^. While KRAS mutations drive a decoupling of glucose and glutamine metabolism in cancer cells, this supports their growth ^29^. While a link between KRAS and LAT2 has not been reported, increased demand for glutamine in KRAS-mutant cells (like KP4) may upregulate LAT2 or similar transporters. FLRT2 is a member of the FLRT (Fibronectin-Leucine Rich Transmembrane) family. It is involved in cell adhesion and guidance and plays a role in metastasis ^30^. There is also currently no published link between KRAS and FLRT2; however, both KRAS and FLRT2 are plasma membrane-associated proteins: KRAS anchors to the inner leaflet via lipid modification, while FLRT2 is a transmembrane adhesion receptor with extracellular and intracellular domains.

SEMA4C, a semaphorin family member, is reported to activate Rho GTPase and PI3K/AKT signalling to drive EMT and invasive behaviour ^31^. While not yet studied extensively in KRAS-mutant PDAC, semaphorins including SEMA4C, have been implicated in pancreatic tumour progression and metastatic spread ^32^. SEMA4C is also a type I transmembrane protein localised at the plasma membrane, which may explain its proximity to KRAS.

### Two dataset analysis

The same 18 raw files were searched using FragPipe (Version 23.1, Uniprot Release 2025_08) as described in the methods section, and the resulting *combined_protein.tsv* was used for the two-dataset analysis. The app was switched to “Two dataset comparison mode” and the same MaxQuant *proteinGroups* data was uploaded, along with its experimental template. On clicking the “Two dataset comparison mode” a second pair of file input boxes appears, the “Data File type 2” was selected as FragPipe, using the default “LFQ” data type and default “MaxLFQ Intensity”. A new experimental template was generated, and this was completed using a spreadsheet editor, to contain the same condition names as the MaxQuant Data. Finally, a comparison file was uploaded that generated the comparisons KRas_KP4 vs Con_KP4, KRas_Suit2 vs Con_Suit2, KRas_h6c7 vs Con_h6c7, for both datasets. This was identical to the 1^st^ round comparisons used is the previously described results and represents the proteins intensities that were enriched over the matched BirA only control. As before, MaxQuant data contained 2795 proteins groups of which 1939 (69%) proteins were reproducibly quantified after filtering. The FragPipe data contained 3504 protein groups, of which 3267 (93%) were reproducibly quantified after filtering. This FragPipe data contained significantly more proteins after filtering suggesting that there were fewer missing values. When the protein numbers per sample were inspected, and the values were downloaded it could be seen that MaxQuant samples has an average of 1315 proteins per sample, but the FragPipe data had an average of 2309 proteins per sample (Supporting Table 1). Both fold-change cut-off values were initially set to >= 10 (Log_2_ value 3.32) for each comparison, with identical adjusted p-value cut-off values of <= 0.05 this resulted in the same 192 significant proteins for the MaxQuant data and 142 proteins for the FragPipe data and an overlap of 72 proteins. To make the significant protein number to be more comparable, the fold-change cut-off value for FragPipe data was lowered to 8 (Log_2_ value 3), which gave 184 significant proteins and a larger overlap of 86 proteins.

### Two data cluster analysis

When the resulting comparison heatmaps were inspected (Supporting Figures S5A and S5B), the FragPipe data had less contrast in the protein intensities between the BirA_KRAS samples and the BirA control, as indicated in a less deep red heatmap values, especially in Cluster 4 which mainly contained proteins that were significantly changed in more than one comparison. The lower contrast may be due to the lower number of missing values in the FragPipe data, which is indicated in higher amount of protein per sample. The second feature of the comparison heatmaps is the low amount of overlap between the two datasets, with only 86 (29.7%) of the significant proteins overlapping (Figure 5A, Supporting Figure S5C). Further inspection of the comparison heatmaps showed very little overlap in Cluster 1, which represents proteins that have higher intensities in the control samples and are not specific to KRAS interactions. Cluster 4 had the most overlapping significant proteins, which also represents proteins that had the highest intensities compared with the control samples. However, there were 4 proteins (SPTBM2, TP53, DLG1, PVRL2) which were significantly changed in all three comparisons in the MaxQuant data but were not seen in the FragPipe data (Figures 5B-C). SPTBM2 and TP53 or any of its aliases were not found in the FragPipe data set. DLG1 was in the FragPipe data set but had a maximum Log_2_ value of 2.62 so was excluded. PVLR2, however, is the alias of NECTIN2, which was also significant in all three comparisons for the FragPipe data. RAB6B, THSD1 and ABCC5 were significant in all three comparisons for the FragPipe data but not significant for the MaxQuant data. RAB6B has 90% identity with RAB6A, which is in the MaxQuant dataset but not significant. ABCC5 was in the MaxQuant dataset but was not significant. However, neither THSD1 nor its aliases were in the MaxQuant dataset.

**Figure 5.**
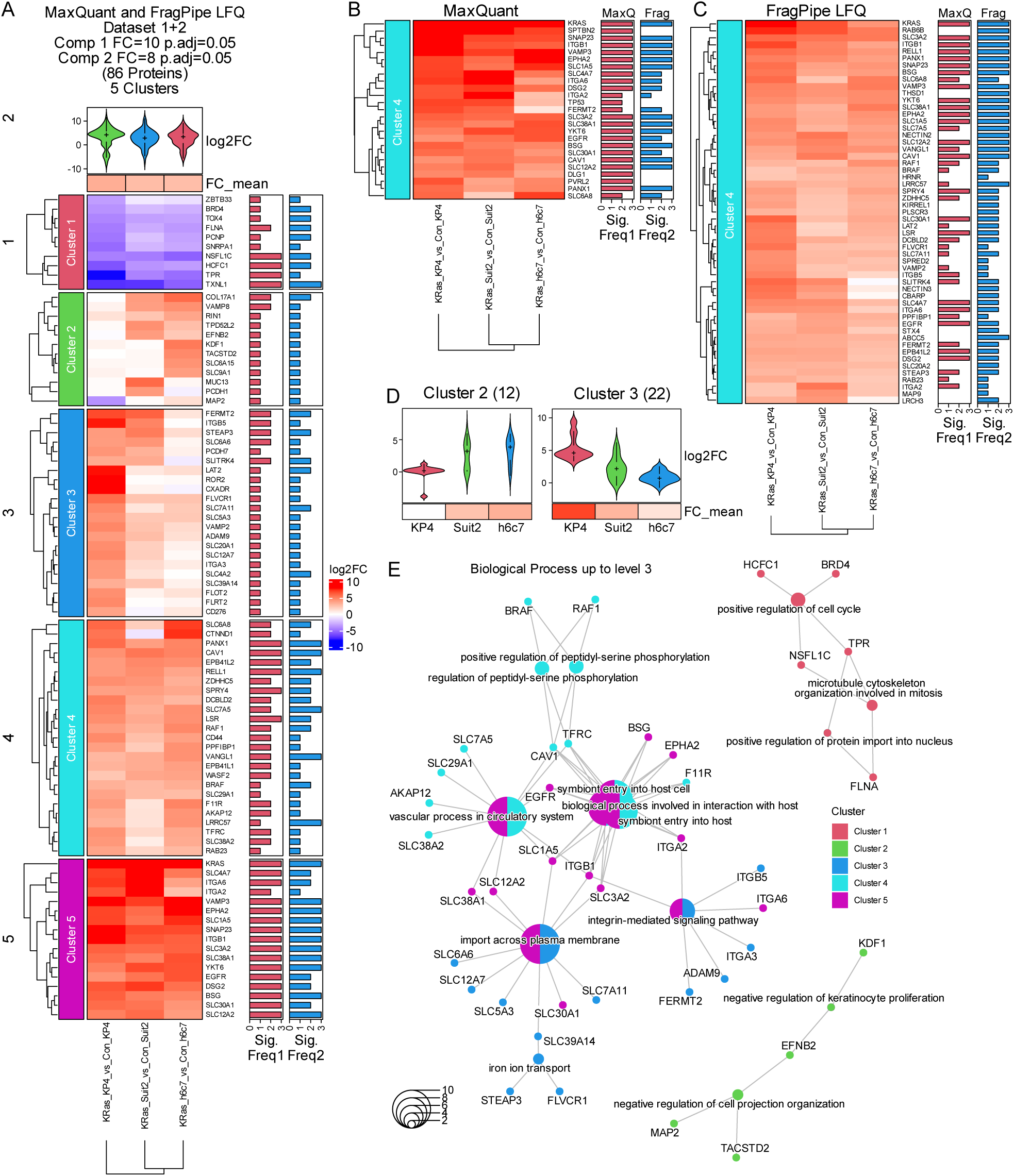
Comparison of MaxQuant and FragPipe datasets. **(A)** Heatmap displays log₂ fold-changes of 86 differentially expressed proteins (FC ≥ 10 or 8, adj. p ≤ 0.05) that were significant for both datasets using comparisons KRas_KP4 vs Con_KP4, KRas_Suit2 vs Con_Suit2, KRas_h6c7 vs Con_h6c7 (Supplementary Figure S5A). **(B-C)** Partial Heatmaps, showing Cluster 4 proteins only for the single dataset heatmaps for MaxQuant and FragPipe. **(D)** Violin plot and means for log₂FC values for Clusters 2 and 3 of the overlap heatmap (panel A). **(E)** Category-gene network plot shows enriched Biological Process GO terms in Clusters 1-5 of the overlapping 86 significant proteins

The 86 overlapping proteins could be separated out into 5 distinct clusters (Figure 5A). Cluster 1 was discounted as it contains proteins that are enriched in the control samples. Cluster 2 contains proteins with higher intensities in Suit2 and H6c7 cells and Cluster 3 higher intensities in KP4 cells (Figure 5D). Clusters 4 and 5 represent proteins enriched in the KRAS samples for all three cell lines. When, the 86 overlapping proteins were investigated, however, many were found to map to similar biological processes. Twenty of these proteins belong to the Solute Carrier (SLC) family (including LAT2), suggesting that KRAS may interact with or be spatially associated with this class of proteins. Gene Ontology analysis revealed that 38 of the overlapping proteins form a coherent Biological Process network (Figure 5E). The main bulk of the network consists of proteins from Cluster 3, 4, and 5. Proteins with higher intensities in KP4 cells (Cluster 3) predominantly mapped to importation across the plasma membrane (symporters) and integrin-mediated signalling, which matches the conclusions of the MaxQuant data cell line comparison. Proteins in Cluster 4 and 5, which are KRAS interactors common in all cell lines, also matched to “Symbiont Entry into Host Cell”, which includes attachment and membrane fusion, and “vascular process in circulatory system”, which describes the development, maintenance, and regulation of blood vessels and vascular function within the circulatory system. When the cellular component was investigated, all the proteins were found to be in or near the basal plasma membrane, with a majority forming parts of focal adhesions (Supporting Figure S3E), which matched the MaxQuant comparisons (Figure 3B).

### Conclusion for two dataset analysis

The two-dataset analysis of the MaxQuant and FragPipe revealed a strong enrichment of Solute Carrier (SLC) family proteins within the KRAS-associated interactome identified by BioID. Notably, SLC1A5 (ASCT2), SLC3A2 (CD98), and SLC7A5 (LAT1), which form a functional axis that regulates amino acid transport and mTORC1 activation. SLC1A5 mediates glutamine uptake, which is subsequently exchanged for leucine via the LAT1/CD98 complex, a process essential for mTORC1 signalling and cellular growth ^33^. Among the 25 most abundant proteins from the overlapped significant protein list, all three proteins were in the top 10 in both datasets (Figure 6A and B). In another KRAS BioID study, SNAT1 (SLC38A1) together with SLC1A5 and SLC7A5, which are among the most abundant significant proteins, were enriched in TurboID-KRAS_G12D-expressing HEK293T cells, compared with wild-type and G12C and G12V mutants ^34^. LAT2 (SLC7A8), which was significant in the cell type comparison and had a greater protein intensity in the KP4 cells, also partners with CD98 ^35^., suggesting this plays a greater role in mTORC1 signalling in that cell type.

**Figure 6.**
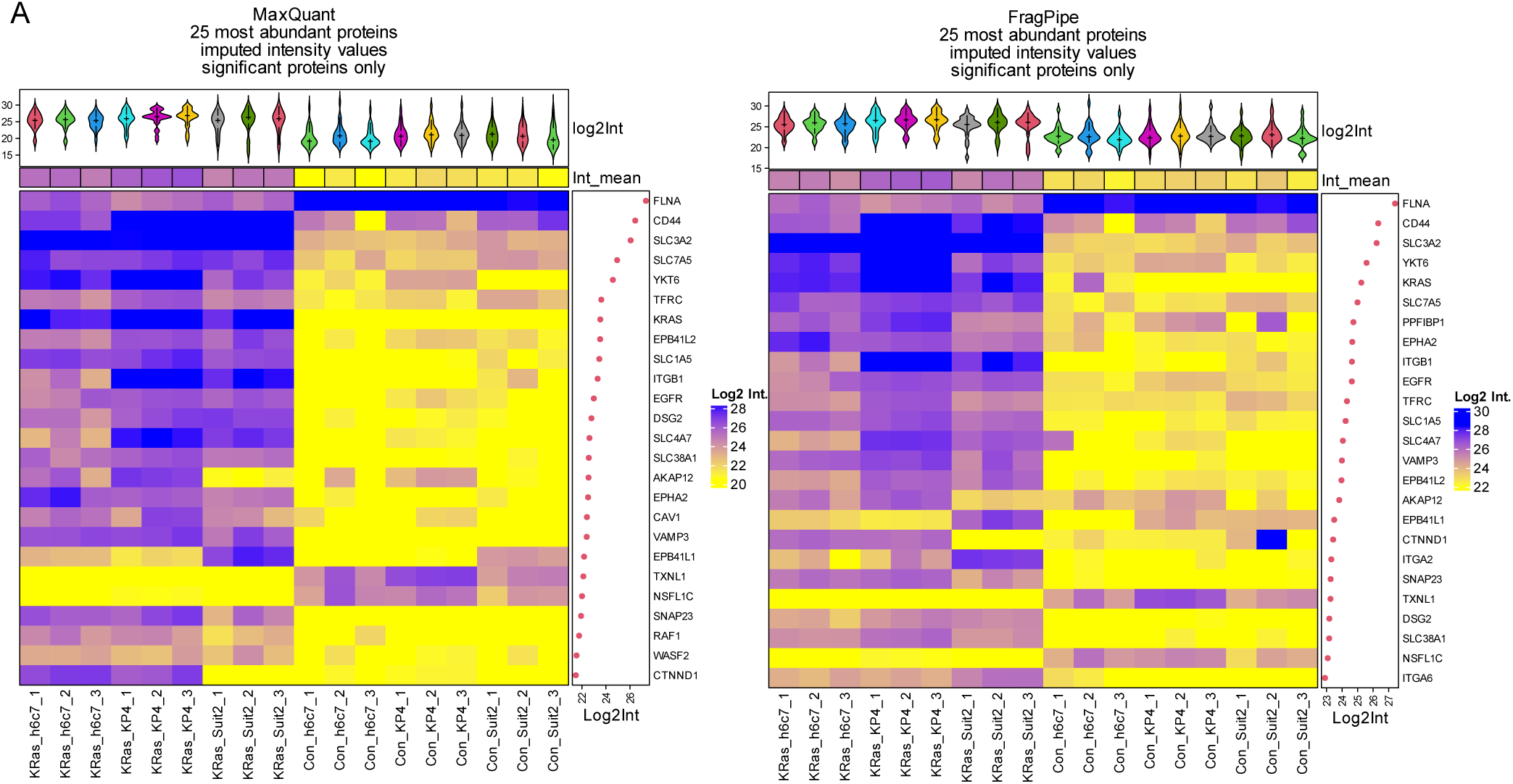
Abundance heatmaps for significant proteins. Heatmaps generated by Manchester Proteome Profiler showing the log₂ intensity values of the 25 most abundant proteins from the significant protein list of 86 proteins from the **(A)** MaxQuant and **(B)** FragPipe datasets. The proteins are ranked on their mean intensity across all samples, with the mean intensity shown on the right annotation. The top annotation shows a violin plots showing the intensity distribution for each sample, along with the sample mean value. The number of proteins shown in the abundance heatmaps can chosen in the app, and the corresponding datable can be downloaded.

Beyond amino acid transport, SLC4A7, which is also in the top 25 abundant proteins, has been implicated in nucleotide biosynthesis downstream of mTORC1 ^36^, while variants of CD44, which is the second most abundant protein in both datasets, stabilises the xCT antiporter (SLC7A11/SLC3A2), promoting cysteine uptake and redox balance ^37^. Other candidates in the top 25 most significant proteins included trafficking proteins YKT6, VAMP3, and SNAP23, which were identified as RAS interactors in a previous BioID study, suggesting a role for vesicular transport in KRAS-driven signalling ^38^. This study also identified the relationship between SLC3A2 and RAS in multiple myeloma, which was further established when a BioID2_SLC3A2 construct also showed KRAS as an interactor.

Several amino acid transporters, including SLC1A5, SLC7A5, SLC7A11, and SLC3A2, are frequently upregulated in cancer, underscoring their role in metabolic reprogramming ^39^. SLC1A5 has emerged as a critical downstream effector of KRAS signalling, with elevated expression in KRAS-mutant colorectal cancer ^40^. However, functional studies suggest ASCT2 (SLC3A2) primarily acts as an exchanger rather than a net importer, with compensatory mechanisms via SNAT transporters, SLC38A1 (SNAT1) and SLC38A2 (SNAT2) maintaining glutamine uptake upon ASCT2 loss ^41^. The SNAT transporters are highly enriched in the KRAS BioID samples, showing their proximity to KRAS. Collectively, these findings highlight a complex network of nutrient transporters and trafficking proteins that integrate metabolic and signalling pathways downstream of KRAS, offering potential therapeutic targets in cancer.

## Conclusion

Manchester Proteome Profiler is a versatile tool for downstream proteomic analysis. It can import grouped protein intensity data from multiple sources. Coupled with a simple file linking the column titles with the experimental conditions, and a further simple file stipulating which experimental conditions should be compared, the data can be transformed into a rich information knowledge tool. Users can quickly assess the quality of the data, reproducibility among replicates and the number and abundance of the proteins. The second part of the app allows users to analyse how the protein intensities of their different experimental conditions differ from each other. The data comparisons can be rapidly changed by uploading a different comparison table. The user can then interrogate how the results of the comparisons cluster into different groups. These clusters can then be further analysed using GO analysis, STRING or BioGRID to identify themes or interaction networks. Researchers have a variety of choices to analyse their raw data, be it open-source packages such as MaxQuant or FragPipe, or commercial packages such as Proteome Discoverer or Spectronaut. Manchester Proteome Profiler allows users to compare the data from different analysis packages to give a more cohesive robust target list for their downstream research. Results of mass spectrometry studies should be followed by validation of the candidate proteins. However, follow up experiments can be costly in both time and research costs, so allowing the researcher to thoroughly investigate their mass spectrometry results in multiple ways, including original the raw intensities, will facilitate any research project.

A secure public server of Manchester Proteome Profiler is available at https://mpp.sbs.manchester.ac.uk/. Source code will be made available upon peer-reviewed publication.

The dataset files can be found at https://doi.org/10.5281/zenodo.17258450.

## Acknowledgements

This work has been funded by Cancer Research UK project No. DRCRPG-May21\100002. We would like to The University of Manchester Research IT and Manchester Cell-Matrix Centre for their help setting up the public server.

