## Supporting Data for "Manchester Proteome Profiler: A User-Friendly Platform for Quantitative Proteomic Analysis"

### Supporting Material

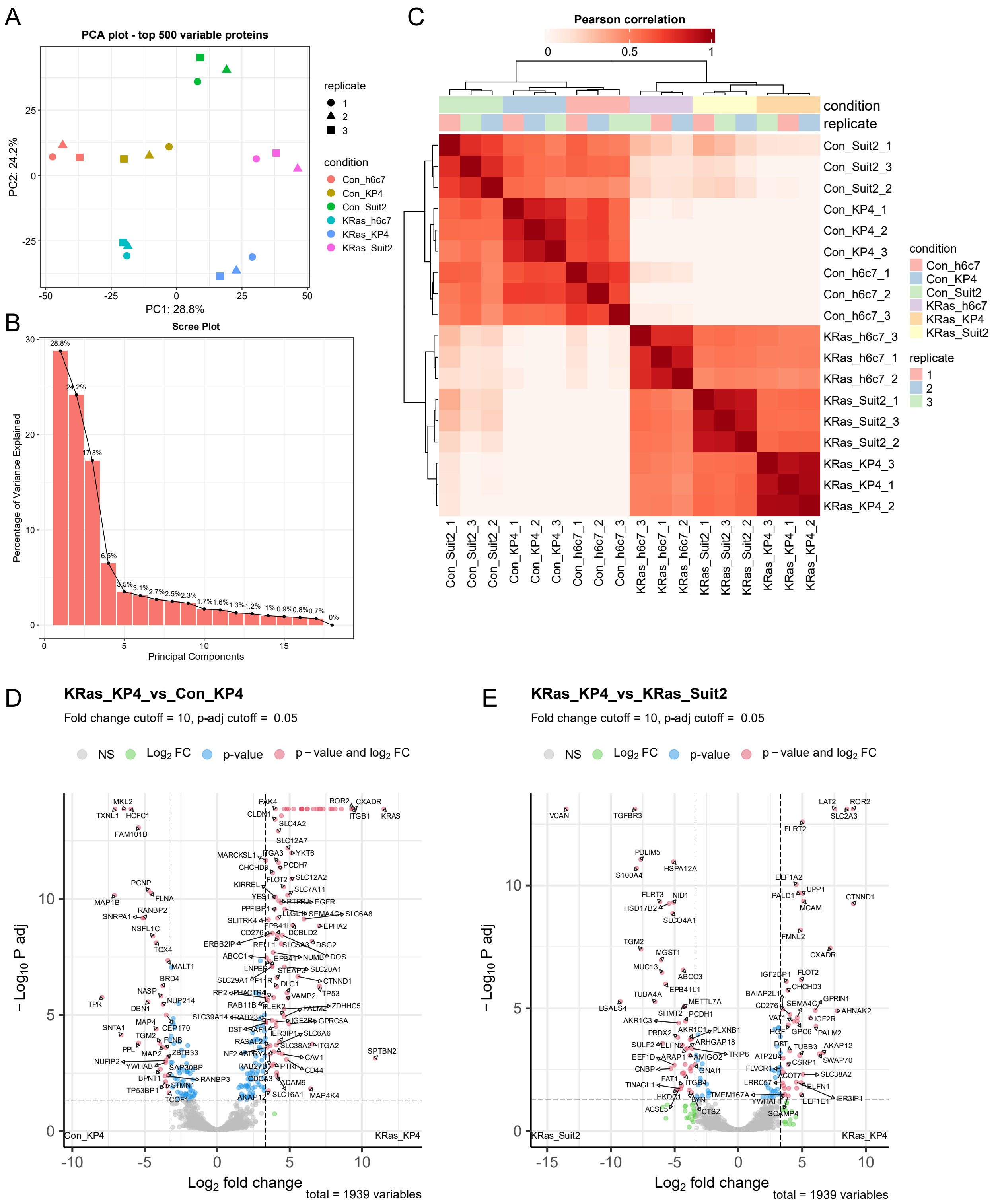


**Supporting Figure S1.** Quality Control and initial results plots for Manchester Proteomic Profiler. **(A)** Principal Component Analysis (PCA) of the top 500 most variable proteins across samples. Each point represents a sample, coloured by condition and shaped by replicate number. **(B)** Scree plot showing the proportion of variance explained by the first 20 principal components. **(C)** Heatmap ordered by hierarchical clustering based on Pearson correlation of protein expression profiles across samples. Blocks of high correlation are shown in dark red. **(D)** A representative volcano plot comparing protein intensities from the 1st round of comparisons KRas_KP4 vs Con_KP4 cells. Plotted is the $-\log_{10}$ P.adj values verses $\text{Log}_2$ fold-change and proteins are labelled if they pass the cutoff values indicated by the dotted lines. $-\log_{10}$ P.adj rather than $-\log_{10}$ P-values is preferred to give a better representation of the data. This results in the volcano plates having a U shape rather than the classical V shape seen with $-\log_{10}$ P-values plots. **(E)** A representative volcano plot from the second set of comparisons KRas_KP4 vs KRas_Suit2.

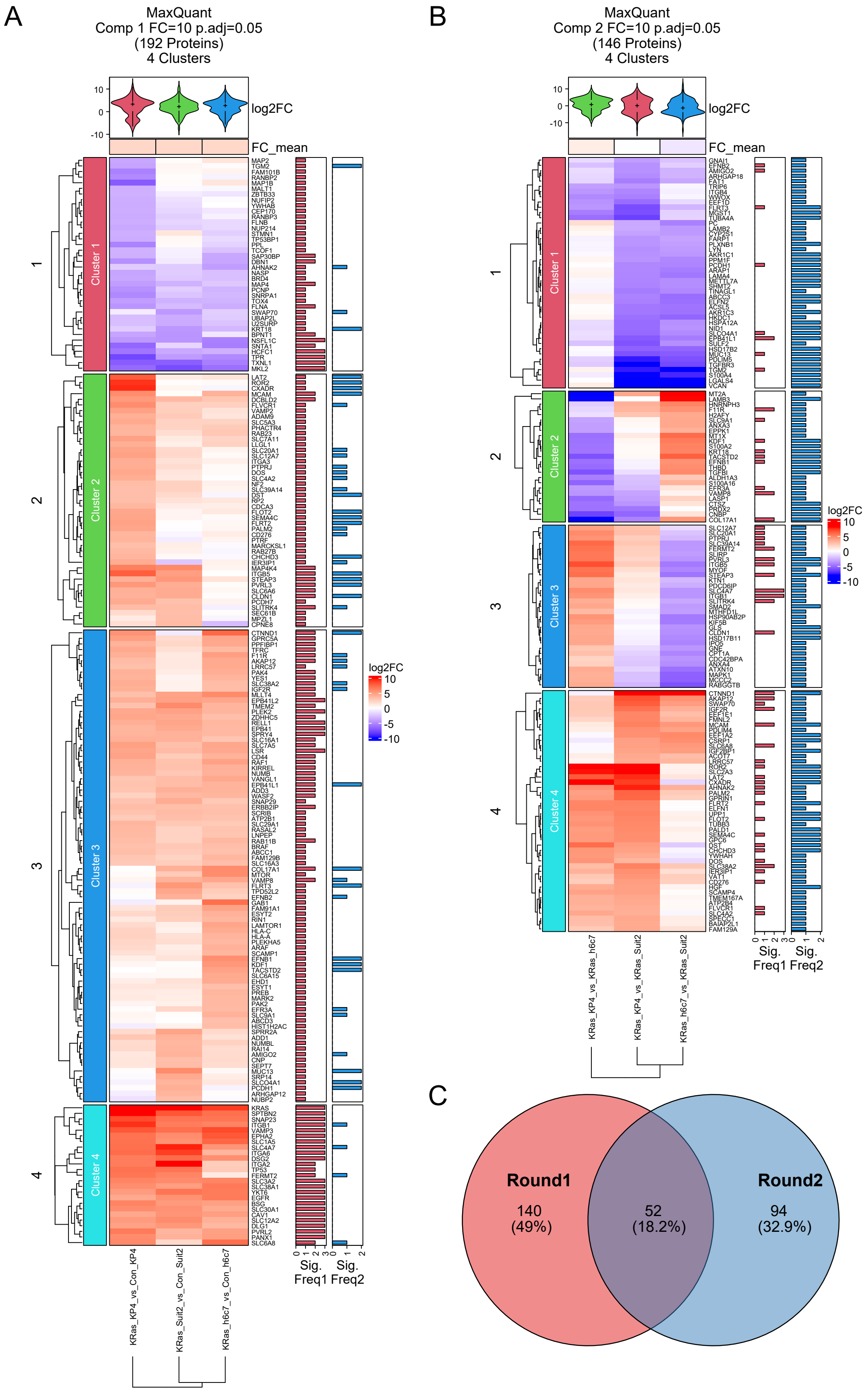


**Supporting Figure S2.** Heatmap and volcano plots generated by Manchester Proteome Profiler. **(A)** Heatmap displays $\log_2$ fold-changes of 192 differentially expressed proteins ($FC \geq 10$, adj. $p \leq 0.05$) that were significant for the comparisons KRas_KP4 vs Con_KP4, KRas_Suit2 vs Con_Suit2, KRas_h6c7 vs Con_h6c7. **(B)** Heatmap displays $\log_2$ fold changes of 146 differentially expressed proteins ($FC \geq 10$, adj. $p \leq 0.05$) that were significant for the comparisons KRas_KP4 vs KRas_Suit2, KRas_KP4 vs KRas_h6c7, KRas_h6c7 vs KRas_Suit2. For both heatmaps, k-means clustering grouped the proteins into four clusters, shown along the left. The violin plot summarises the distribution of fold changes for each comparison and the bar plots to the right indicate the number of times the corresponding protein was significant for the first set of comparisons (Sig. Freq 1) and the second set of the comparisons (Sig. Freq 2). **(C)** Venn diagram showing the overlap of the 192 significant proteins from comparison set 1 (Round 1) and the 146 significant proteins from comparison set 2 (Round 2), resulting in the 52 proteins used for subsequent analysis.

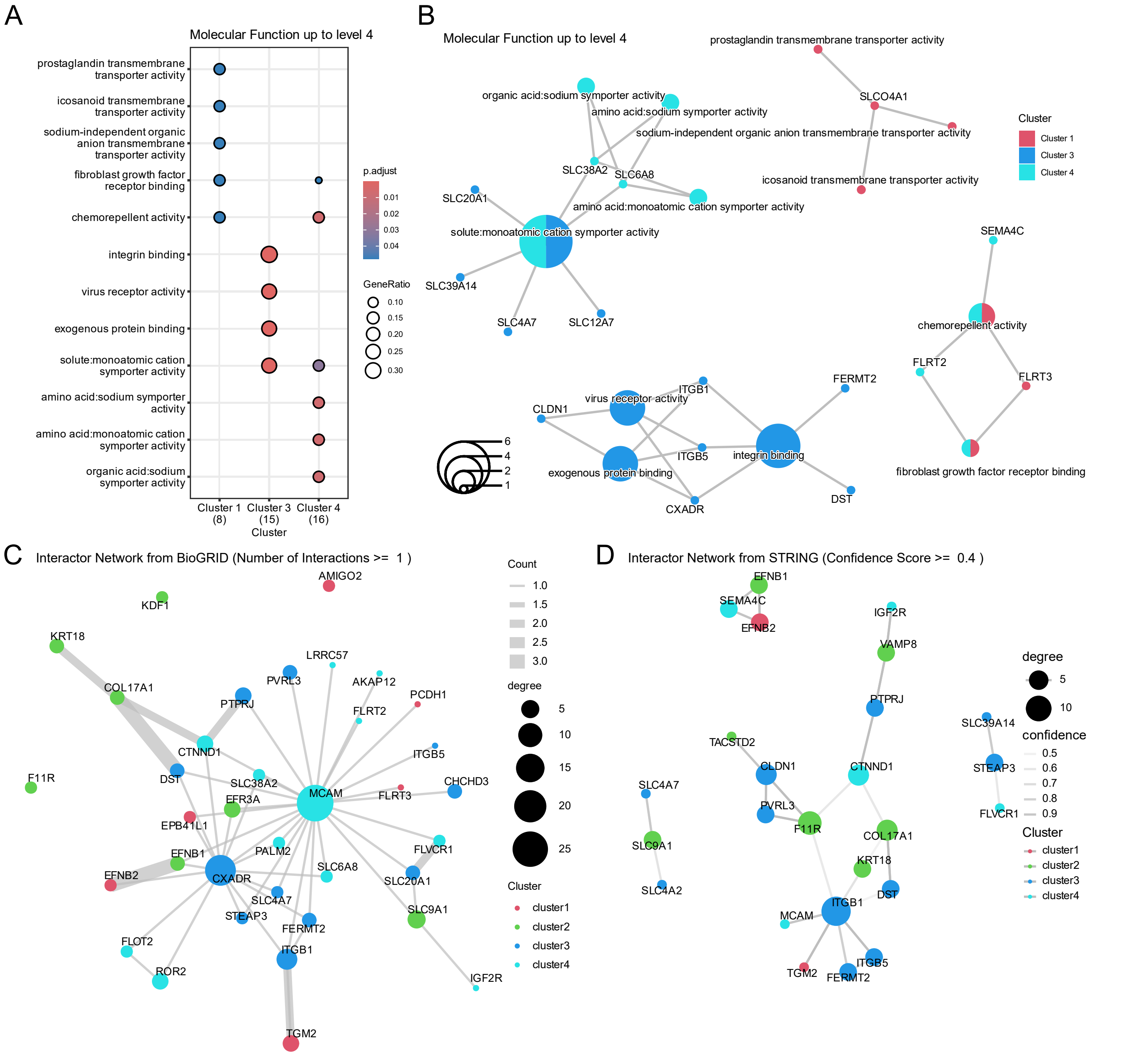


**Supporting Figure S3.** Gene Ontology, BioGRID and STRING plots generated by Manchester Proteome Profiler. **(A)** Dot plot shows enriched Molecular Function GO terms in Clusters 1, 3 and 4. Cluster 2 showed no enriched GO terms. **(B)** Corresponding proteins from each Molecular Function GO term shown as a category-gene network plot (cnetplot). Node colours correspond to the same cluster colours as the heatmap. **(C)** Network plot generated from interactions found from a BioGRID search, a pair of proteins must have at least 1 reported interaction, to be shown and the width of the connecting line is proportional to the number of interactions in the database. Proteins shown with no connecting line indicate self-interactions. The size of the node is proportional to the number of connecting proteins and the colour of the node is representative of the Cluster number. **(D)** Network plot generated from interactions found from a STRING database search, a pair of proteins must have reported interaction confidence score of 0.4, to be shown and the intensity of the connecting line is proportional to the confidence score. Node size and colour is the same as the Biogrid network plot.

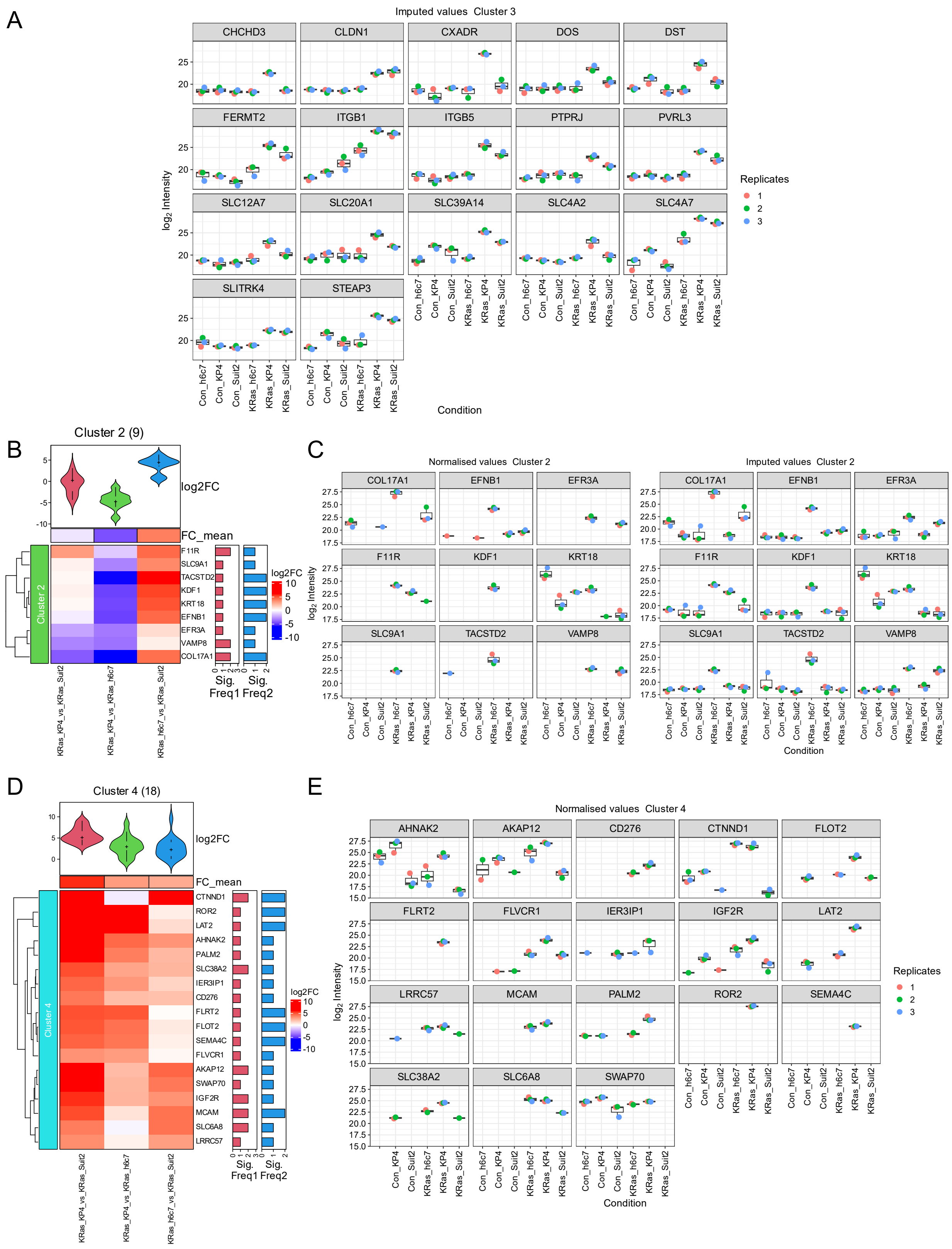


**Supporting Figure S4.** **(A)** Box plots of the imputed protein intensities (using “man” method) of Cluster 3 proteins. The original normalised values are shown in Figure 4B. **(B)** Heatmap of significant proteins in Cluster 2 which represents proteins that are mainly more abundant in Kras_h6c7 samples. **(C)** Box plots showing normalised protein intensities and the imputed protein intensities of the nine proteins in Cluster 2. **(D)** Heatmap of significant proteins in Cluster 4 which represents proteins that are mainly more abundant in Kras_KP4 samples. **(E)** Box plots showing normalised protein intensities of the 18 proteins in Cluster 4. Significant proteins that were only detected in Kras_KP4 samples included FLRT2, ROR2, SEMA4C.

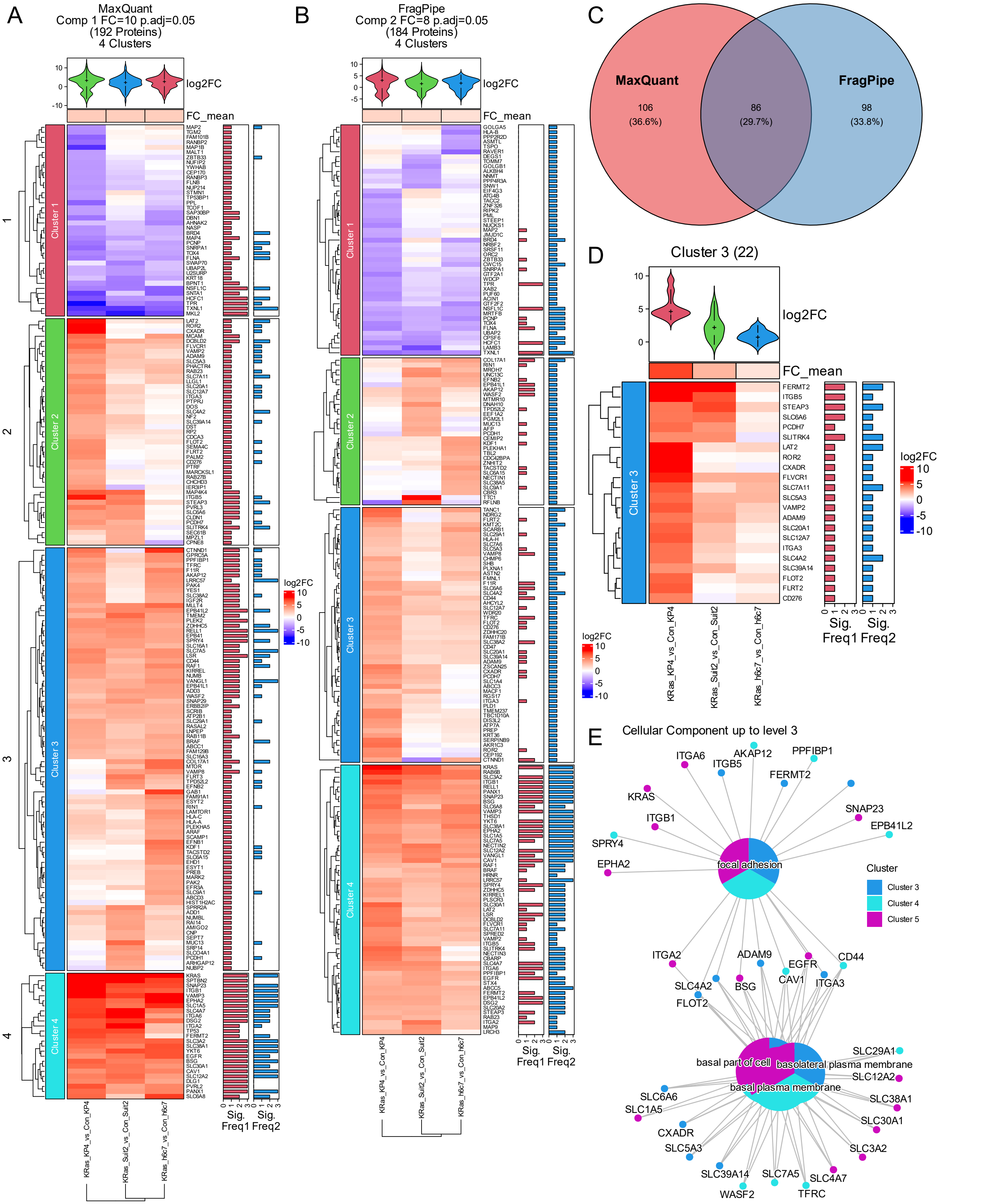


**Supporting Figure S5.** Heatmaps and Venn diagram of two dataset analysis. Heatmap displays $\log_2$ fold changes of differentially expressed proteins that were significant for the comparisons KRas_KP4 vs Con_KP4, KRas_Suit2 vs Con_Suit2, KRas_h6c7 vs Con_h6c7 for the **(A)** MaxQuant dataset ($FC \geq 10$, adj. $p \leq 0.05$) and the **(B)** FragPipe dataset ($FC \geq 8$, adj. $p \leq 0.05$)**.** **(C)** Venn diagram showing the overlap significant proteins from MaxQuant dataset and the FragPipe dataset resulting in the 86 proteins used for subsequent analysis. **(D)** Heatmap of overlapping significant proteins for Cluster 3, which represents proteins that are mainly more abundant in Kras_KP4 samples. **(E)** Biological Process GO terms shown as a category-gene network plot for the overlap heatmap.

**MaxQuant**

| sample | total_proteins | proteins_in_sample | label | condition | replicate |
|---|---|---|---|---|---|
| Con_KP4_1 | 1939 | 1509 | Con_BirA_ | Con_KP4 | 1 |
| Con_KP4_2 | 1939 | 1494 | Con_BirA_ | Con_KP4 | 2 |
| Con_KP4_3 | 1939 | 1419 | Con_BirA_ | Con_KP4 | 3 |
| Con_Suit2_1 | 1939 | 1485 | Con_BirA_ | Con_Suit2 | 1 |
| Con_Suit2_2 | 1939 | 1414 | Con_BirA_ | Con_Suit2 | 2 |
| Con_Suit2_3 | 1939 | 1348 | Con_BirA_ | Con_Suit2 | 3 |
| Con_h6c7_1 | 1939 | 1140 | Con_BirA_ | Con_h6c7 | 1 |
| Con_h6c7_2 | 1939 | 1027 | Con_BirA_ | Con_h6c7 | 2 |
| Con_h6c7_3 | 1939 | 975 | Con_BirA_ | Con_h6c7 | 3 |
| KRas_KP4_1 | 1939 | 1575 | BirA_KRas_ | KRas_KP4 | 1 |
| KRas_KP4_2 | 1939 | 1487 | BirA_KRas_ | KRas_KP4 | 2 |
| KRas_KP4_3 | 1939 | 1384 | BirA_KRas_ | KRas_KP4 | 3 |
| KRas_Suit2_1 | 1939 | 1418 | BirA_KRas_ | KRas_Suit2 | 1 |
| KRas_Suit2_2 | 1939 | 1396 | BirA_KRas_ | KRas_Suit2 | 2 |
| KRas_Suit2_3 | 1939 | 1358 | BirA_KRas_ | KRas_Suit2 | 3 |
| KRas_h6c7_1 | 1939 | 1239 | BirA_KRas_ | KRas_h6c7 | 1 |
| KRas_h6c7_2 | 1939 | 976 | BirA_KRas_ | KRas_h6c7 | 2 |
| KRas_h6c7_3 | 1939 | 1029 | BirA_KRas_ | KRas_h6c7 | 3 |
| | **Average** | 1315 | | | |

**FragPipe**

| sample | total_proteins | proteins_in_sample | label | condition | replicate |
|---|---|---|---|---|---|
| Con_KP4_1 | 3267 | 2533 | Con_BirA_ | Con_KP4 | 1 |
| Con_KP4_2 | 3267 | 2456 | Con_BirA_ | Con_KP4 | 2 |
| Con_KP4_3 | 3267 | 2396 | Con_BirA_ | Con_KP4 | 3 |
| Con_Suit2_1 | 3267 | 2538 | Con_BirA_ | Con_Suit2 | 1 |
| Con_Suit2_2 | 3267 | 2480 | Con_BirA_ | Con_Suit2 | 2 |
| Con_Suit2_3 | 3267 | 2361 | Con_BirA_ | Con_Suit2 | 3 |
| Con_h6c7_1 | 3267 | 2090 | Con_BirA_ | Con_h6c7 | 1 |
| Con_h6c7_2 | 3267 | 1905 | Con_BirA_ | Con_h6c7 | 2 |
| Con_h6c7_3 | 3267 | 1952 | Con_BirA_ | Con_h6c7 | 3 |
| KRas_KP4_1 | 3267 | 2626 | BirA_KRas_ | KRas_KP4 | 1 |
| KRas_KP4_2 | 3267 | 2532 | BirA_KRas_ | KRas_KP4 | 2 |
| KRas_KP4_3 | 3267 | 2370 | BirA_KRas_ | KRas_KP4 | 3 |
| KRas_Suit2_1 | 3267 | 2453 | BirA_KRas_ | KRas_Suit2 | 1 |
| KRas_Suit2_2 | 3267 | 2380 | BirA_KRas_ | KRas_Suit2 | 2 |
| KRas_Suit2_3 | 3267 | 2343 | BirA_KRas_ | KRas_Suit2 | 3 |
| KRas_h6c7_1 | 3267 | 2166 | BirA_KRas_ | KRas_h6c7 | 1 |
| KRas_h6c7_2 | 3267 | 1922 | BirA_KRas_ | KRas_h6c7 | 2 |
| KRas_h6c7_3 | 3267 | 2062 | BirA_KRas_ | KRas_h6c7 | 3 |
| | **Average** | 2309 | | | |

**Supporting Table 1.** Protein Groups per sample for MaxQuant and FragPipe datasets
